# Suv39h-catalysed H3K9me3 is critical for euchromatic genome organisation and the maintenance of gene transcription

**DOI:** 10.1101/2020.08.13.249078

**Authors:** Christine R. Keenan, Hannah D. Coughlan, Nadia Iannarella, Timothy M. Johanson, Wing Fuk Chan, Alexandra L. Garnham, Gordon K. Smyth, Rhys S. Allan

## Abstract

H3K9me3-dependent heterochromatin is critical for the silencing of repeat-rich pericentromeric regions and also has key roles in repressing lineage-inappropriate protein-coding genes in differentiation and development. Here, we investigate the molecular consequences of heterochromatin loss in cells deficient in both Suv39h1 and Suv39h2 (Suv39DKO), the major mammalian histone methyltransferase enzymes that catalyse heterochromatic H3K9me3 deposition. Unexpectedly, we reveal a predominant repression of protein-coding genes in Suv39DKO cells, with these differentially expressed genes principally in euchromatic (DNaseI-accessible, H3K27ac-marked) rather than heterochromatic (H3K9me3-marked) regions. Examination of the 3D nucleome reveals that transcriptomic dysregulation occurs in euchromatic regions close to the nuclear periphery in 3-dimensional space. Moreover, this transcriptomic dysregulation is highly correlated with altered 3-dimensional genome organization in Suv39DKO cells. Together, our results suggest that the nuclear lamina-tethering of Suv39-dependent H3K9me3 domains provides an essential scaffold to support euchromatic genome organisation and the maintenance of gene transcription for healthy cellular function.

## Introduction

Gene silencing in regions of heterochromatin is critical to cell identity and appropriate cell-fate decisions during development and differentiation (Nicetto et al., 2019; Yadav et al., 2018). Within all eukaryotic nuclei, heterochromatin and euchromatin are spatially segregated, with euchromatin typically located in the nuclear interior and heterochromatin at the nuclear periphery in large lamina-associated domains (LADs) often spanning several megabases in size (van Steensel and Belmont, 2017). The mechanisms driving the formation of these distinct compartments have been unclear but recent modelling suggests that biophysical attractions between heterochromatic regions, and not euchromatic regions, drives the phase separation of active and inactive domains leading to the subsequent tethering of heterochromatic domains to the nuclear-lamina (Falk et al., 2019).

In mammalian cells, the heterochromatin-associated mark H3K9me3 is deposited by the suppressor of variegation 3-9 homologue (Suv39h) enzymes Suv39h1 and Suv39h2. This H3K9me3 histone mark is then recognised by heterochromatin protein-1 (HP1)-corepressor reader molecules that self-oligomerise to impart gene silencing and facilitate the tethering of heterochromatin domains to the nuclear lamina (Bannister et al., 2001; Lachner et al., 2001; Towbin et al., 2012). H3K9me3-dependent heterochromatin is critical for the silencing of repeat-rich pericentromeric regions (Bulut-Karslioglu et al., 2014) and also has key roles in repressing lineage-inappropriate genes (Allan et al., 2012; Liu et al., 2015; Nicetto et al., 2019) and impeding the binding of diverse transcription factors and RNA polymerase, unlike H3K27me3-marked domains (Becker et al., 2017; Becker et al., 2016).

Of note, loss of heterochromatin has been proposed as a potential universal molecular cause of ageing, whereby over time, heterochromatin domains lose integrity leading to de-repression of silenced genes and thus aberrant expression patterns and cellular dysfunction (Tsurumi and Li, 2012; Villeponteau, 1997). Loss of function of the immune system is a key feature of ageing whereby elderly individuals become less responsive to vaccination and more susceptible to a range of infections (Keenan and Allan, 2019). Mice deficient in both Suv39h1 and Suv39h2 enzymes (Suv39DKO) do not survive embryonic development on a pure C57Bl/6 background (Keenan et al., 2020), unlike those on a mixed background where some developmentally defective mice can survive to birth at sub-mendelian ratios (Peters et al., 2001). We have recently developed a chimeric model in which we can reconstitute the entire haematopoietic compartment of donor mice with Suv39DKO donor stem cells to examine the premature ageing of the Suv39DKO immune system (Keenan et al., 2020), providing us now with an opportunity to elucidate the molecular mechanisms of cellular dysfunction following heterochromatin loss.

Here, we investigate the molecular consequences of loss of heterochromatin in Suv39DKO primary immune cells. We unexpectedly reveal predominant gene repression, not activation, following heterochromatin loss, with differentially expressed genes principally in euchromatic regions rather than heterochromatic regions. We explore the mechanism of this altered transcriptional profile by examining the 3D nucleome through LaminB1 ChIP-seq and *in situ* Hi-C. Together, our results suggest that the nuclear lamina-tethering of Suv39-dependent H3K9me3 domains provides an essential scaffold to support euchromatic genome organisation and the maintenance of gene transcription for healthy cellular function.

## Methods

### Mice

*Suv39h1* and *Suv39h2* null mice (a generous gift from Thomas Jenuwein, Max Planck Institute, Frieburg) were backcrossed to the C57BL/6 strain on which background the Suv39 double-deficient (DKO) genotype was found to be embryonic lethal (Keenan et al., 2020). We therefore used 8 – 12 week old chimeric mice for this study as previously described (Keenan et al., 2020). Briefly, chimeric mice were generated using *Suv39h1* and *Suv39h2* double-deficient (DKO) CD45.2^+^ foetal liver or bone marrow donor cells which were injected into the tail vein of lethally irradiated (2x 550 *Gy*) congenic (CD45.1^+^) recipient mice. These DKO cells are compared to littermate control cells (designated CON) of *Suv39h1*^+/y^*Suv39h2*^+/−^ genotype, as we used a mating strategy unable to produce wildtype cells to increase the frequency of the sub-mendelian DKO genotype. Our previous studies have shown no defects in development, function or heterochromatin structure of *Suv39h2*^−/−^ mice making the *Suv39h2*^+/−^ a reasonable surrogate for a true wildtype. All mice were maintained at The Walter and Eliza Hall Institute Animal Facility under specific-pathogen-free conditions. All animal experiments were approved in advance by the Walter and Eliza Hall Institute Animal Ethics Committee and conducted in accordance with published guidelines.

### Flow cytometric sorting

Single cell suspensions were generated from mouse thymus by mechanical homogenization following by red blood cell lysis (156mM NH_4_Cl, 11.9mM NaHCO_3_, 97μM EDTA). Fluorochrome-conjugated antibodies against the following mouse antigens were then used for sorting by flow cytometry: CD45.2-FITC (clone 104) and CD19-PECy7 from BD Pharmingen; CD4-PE (clone GK1.5) and CD8α-APC (clone 53.6.7) which were generated internally. Surface staining was carried out at 4°C for 30 mins. Live (Propidium iodide (ThermoFisher) negative), CD45.2^+^CD19^−^CD4^+^CD8^+^ cells were sorted using BD FACSAria™ cell sorter (***Supp Fig S1***)

### RNA-seq

RNA was extracted using RNeasy Plus Mini Kit (Qiagen) and quantified in a TapeStation 2200 using RNA ScreenTape (Agilent). Libraries for sequencing were prepared using the TruSeq RNA sample-preparation kit (Illumina) from 500 ng RNA, amplified with KAPA HiFi HotStart ReadyMix (Kapa Biosystems), size-selected to 200 – 400 bp, and cleaned up with AMPure XP magnetic beads (Beckman). Final libraries were quantified with TapeStation 2200 using D1000 ScreenTape for sequencing on the Illumina NextSeq 500 platform to produce 81 bp paired-end reads. Around 35 million read pairs were generated per sample.

All read pairs were aligned to the mouse genome, build mm10, using align from the Rsubread package (Liao et al., 2019) v1.24.0. Over 91% of all read pairs were successfully mapped for each sample. The number of read pairs overlapping genes were summarized into counts using Rsubreads featureCounts (Liao et al., 2014). Of those successfully mapped, an average of 89% of these read pairs were assigned to a gene. Genes were identified using Gencode annotation to the mm10 genome, version M22. Differential expression analyses were then undertaken using the limma (Ritchie et al., 2015) and edgeR (Robinson et al., 2010) software packages, versions 3.42.2 and 3.28.1 respectively.

Prior to analysis all genes with no symbol, non-protein coding and Riken genes were removed. Expression based filtering was also performed. All genes were required to achieve a count per million reads (CPM) greater than 0.4 in at least 3 samples to be retained. Following filtering 11,036 genes remained. Compositional differences between samples were then normalized using the trimmed mean of M-values (TMM) method (Robinson and Oshlack, 2010). All counts were transformed to log_2_-CPM with associated precision weights using voom (Law et al., 2014). Differential expression between the knock-out and wild type samples was evaluated using linear models and robust empirical Bayes moderated t-statistics relative to a fold-change threshold of 1.2 (McCarthy and Smyth, 2009; Phipson et al., 2016). P-values were adjusted to control the false discovery rate (FDR) below 5% using the Benjamini and Hochberg method.

Analyses of the Gene Ontology (GO) terms (The Gene Ontology, 2019) was performed using limma’s goana function. Gene set testing of the ageing signature and corresponding barcode plots were generated using limma’s roast and barcodeplot functions. The ageing signature was evaluated using both directional and non-directional roast tests with 9,999 rotations. The heatmap was generated using the pheatmap package available through CRAN.

### ChIP-seq

Cells (>5M) for LaminB1 ChIP-seq were fixed in 1% formaldehyde for 20 min at room temperature with mixing, then quenched by the addition of glycine to a final concentration of 125mM. Fixed cells were washed twice with ice-cold PBS then the cell membrane was lysed by incubation in ChIP buffer (150mM NaCl, 50mM Tris, 5mM EDTA, 0.5%(v/v) NP-40, 1%(v/v) Triton X-100) containing 1x protease inhibitors (Roche) on ice for at least 10 minutes prior to dounce homogenization. Chromatin within intact nuclei was then digested for 5 minutes with micrococcal nuclease (NEB #M0247, 500U/1M cells) prior to nuclei lysis by Covaris sonication. Nuclear lysate was then cleared by centrifugation (>15,000g for 1 min), a small aliquot was taken as a whole cell extract (WCE) control, and LaminB1 antibody (Abcam #ab16048, 3μg) was added to remaining cleared lysate and incubated overnight at 4°C with rotation. Immunoprecipitation was performed using Protein G Dynabeads (ThermoFisher) then chromatin was eluted from beads using elution buffer (1%(v/v) SDS, 0.1M NaHCO_3_). ChIP and WCE samples were then adjusted to 200mM NaCl, digested with RNaseA (20μg/sample) overnight at 65°C, then digested with proteinase K (20μg/sample) for a further 1 h at at 65°C. DNA was then extracted using Zymo ChIP DNA clean and concentrator kit (#D5205) and sequencing libraries were prepared using Illumina TruSeq DNA Sample Prep kit using half volumes of all reagents. Libraries were sequenced on an Illumina NextSeq500 to produce 81 base length paired-end reads. Independent biological duplicates were prepared for each condition and sequenced to a depth of approximately 50 million reads.

### ChIP-seq processing and peak calling

Biological replicate fastq files were combined for ChIP and WCE samples. Fastq files were aligned to the mouse genome (mm10) with Rsubread package v1.28 (Liao et al., 2019). Bam files were sorted with Rsamtools and duplicate reads were removed with MarkDuplicates (Picard suite v1.117 (https://broadinstitute.github.io/picard/). WCE samples were down sampled to the size of the corresponding ChIP libraries (~53 million reads) using DownsampleSam from Picard tools with strategy chained and accuracy 0.001. For viewing and plotting, bedGraph files of the log of the ratio of the ChIP coverage to the WCE coverage were created with bamCoverage from Deeptools (Ramirez et al., 2016) and R. Enriched domain detector (EDD) (Lund, Oldenburg and Collas, 2014) was used to call LADs on the control and DKO samples separately using the WCE as input for the corresponding ChIP. EDD was run ten times without specifying the bin size and gap penalty. The mean of the estimated bin size and gap penalty from the ten trials was used to call LADs (WT: gap penalty 4.79 and bin size 6 kbp, KO: gap penalty 9.1 and bin size 8 kbp). LADs called on the Y and scaffold chromosomes were excluded from subsequent analyses.

### Hi-C

*In situ* Hi-C was performed on fixed cells as previously described (Johanson et al., 2018). Libraries were sequenced on an Illumina NextSeq500 to produce 81 base length paired-end reads. Independent biological duplicates were prepared for each condition and sequenced to a depth of approximately 200 million reads (***Supp Table 1***).

### Hi-C data processing

Data pre-processing and analysis was performed as previously described with changes in parameters (Johanson et al., 2018). Samples were aligned to the mm10 genome using the *diffHic* package v1.14.0 (Lun and Smyth, 2015a) which utilizes cutadapt v0.9.5 (Martin, 2011) and bowtie2 v2.2.5 (Langmead and Salzberg, 2012) for alignment and Picard suite v1.117 (https://broadinstitute.github.io/picard/) for fixing mate pair information and marking duplicate reads. Dangling ends and self-circling artefacts were identified and removed if the pairs of inward-facing or outward-facing reads were separated by less than 1000 bp (inward-facing) or 2000 bp (outward-facing). Read pairs with fragment sizes above 1000 bp were removed. Proportion of reads removed through each processing step is tabulated in **Supp Table 1**.

The HOMER HiC pipeline (Heinz et al., 2010a; Heinz et al., 2010b; Heinz et al., 2018) was also used for HiC analysis. After processing with the *diffHic* pipeline, libraries in HDF5 format were converted to the HiC summary format with R. Then input-tag directions were created for each library with the makeTagDirectory function, with the genome (mm10) and restriction-enzyme-recognition site (GATC) specified. Summed biological-replicate tag directories for each cell type were also created.

To confirm the reproducibility of the libraries the HiCRep R package was utilized to quantify the similarity between all libraries with the stratum adjusted correlation coefficient (SCC) (Yang et al., 2017). For every combination of libraries, the raw contact matrices (50 kbp resolution) of individual chromosomes for each replicate were used to compute the SCC with smoothing parameter 3 and maximum distance considered 5 Mbp. For each pairwise comparison, a median SCC was calculated across all chromosomes (***Supp Fig S2***).

Plaid plots were constructed using the contact matrices and the plotHic function from the Sushi R package (Phanstiel, 2015). The inferno color palette from the *viridis* package (Garnier, 2018) was used. The range of color intensities in each plot was scaled according to the library size of the sample to facilitate comparisons between plots from different samples. DI arcs were plotted with the plotBedpe function of the Sushi package. The z-score shown on the vertical access was calculated as −log10 (P-value).

### Detection of A/B compartments

A/B compartments were identified at a resolution of 100 kbp with the method outlined by Lieberman-Aiden et al.(Lieberman-Aiden et al., 2009) using the HOMER HiC pipeline (Lin et al., 2012). With the summed biological replicate tag directories, the runHiCpca.pl function was used on each library with -res 100000 and the genome (mm10) specified to perform principal component analysis to identify compartments. To identify changes in A/B compartments between libraries, the getHiCcorrDiff.pl function was used to directly calculate the difference in correlation profiles. A region was considered to flip compartment if the difference in correlation profile was <−0.1 and was determined from control cell A/B compartments. ChrY was excluded from the analysis.

### Detecting differential interactions (DIs)

Differential interactions (DIs) between the different libraries were detected using the *diffHic* package v1.16.0 (Lun and Smyth, 2015a). Read pairs were counted into 50 kbp bin pairs for all chromosomes. Bins were discarded if had a count of less than 10, contained blacklisted genomic regions as defined by ENCODE for mm10 (Consortium, 2012) or were within a centromeric or telomeric region. Filtering of bin-pairs was performed using the filterDirect function, where bin pairs were only retained if they had average interaction intensities more than 3-fold higher than the background ligation frequency. The ligation frequency was estimated from the inter-chromosomal bin pairs from a 1 Mbp bin-pair count matrix. Diagonal bin pairs were also removed. The counts were normalized between libraries using a loess-based approach with bin pairs less than 100 kbp from the diagonal normalized separately from other bin pairs. Tests for DIs were performed using the quasi-likelihood (QL) framework (Chen et al., 2016; Lund et al., 2012) of the *edgeR* package v3.26.5 (Robinson et al., 2010). A design matrix was constructed using a one-way layout that specified the group to which each library belonged. A mean-dependent trend was fitted to the negative binomial dispersions with the estimateDisp function. A generalized linear model (GLM) was fitted to the counts for each bin pair (McCarthy et al., 2012), and the QL dispersion was estimated from the GLM deviance with the glmQLFit function. The QL dispersions were then squeezed toward a second mean-dependent trend, using a robust empirical Bayes strategy (Phipson et al., 2016). A p-value was computed against the null hypothesis for each bin pair using the QL F-test. P-values were adjusted for multiple testing using the Benjamini-Hochberg method. A DI was defined as a bin pair with a false discovery rate (FDR) below 5%. DIs adjacent in the interaction space were aggregated into clusters using the diClusters function to produce clustered DIs. DIs were merged into a cluster if they overlapped in the interaction space. The significance threshold for each bin pair was defined such that the cluster-level FDR was controlled at 5%. Cluster statistics were computed using the *csaw* package v1.18.0 (Lun and Smyth, 2016). Overlaps between unclustered bin pairs and genomic intervals were performed using the InteractionSet package (Lun et al., 2016). Differential interactivity for DE genes was calculated from 50 kbp interactions with p-value<0.05 before clustering and multiple testing. For each DE gene, the mean logFC from overlapping interactions was calculated.

### Detecting TAD boundaries

TAD breakpoints were detected in each HiC library with TADbit v0.2.0.5 (Serra et al., 2017). Read pairs were counted into 50 kbp bin pairs (with bin boundaries rounded up to the nearest MboI restriction site). The TADbit tool *find_tad* was run on the raw counts specifying normalized=False. Only TAD boundaries with a score of 7 or higher were included in the results.

### Detecting differential TADs

Differential TAD boundaries (DTB) between groups were detected with the *diffHic* and *edgeR* packages (Lun and Smyth, 2015a) using the approach described in Chapter 8 of the diffHic User’s Guide. This approach adapts the statistical strategy recently described for differential methylation (Chen et al., 2017) to identify DTBs. The strength of a putative TAD boundary was assessed based on the up-stream vs down-stream intensity contrast at that genomic loci, defined as the ratio of up-stream to down-stream HiC reads anchored at that genomic region. edgeR was used to test whether the ratio at each loci significantly increased or decreased in absolute size between groups. This method directly assesses differential boundary strength relative to biological variation without needing to make TADs calls in individual samples. Up-stream and down-stream read counts were determined for 200 kbp genomic regions and for 2 Mbp up- and downstream. Low abundance regions with average log2-counts per million below 2 were removed along with regions in blacklisted regions, telomeres or centromeres. Tests for DTBs were performed using the QL framework of edgeR. Regions with FDR below 0.05 were considered to be DTB.

### Detecting differentially interacting promoters (DIPs)

Differentially interacting promoters were detected using across all libraries with the diffHic package (Lun and Smyth, 2015a) and the method described in (Chan et al., 2019) with alterations in parameters. Gene transcriptional start sites were defined with the mouse GENCODE Gene set annotation (v. M24). Promoter regions (upstream 10kbp and downstream 5kbp) were defined with the promoters function from the GenomicFeatures package v1.36.2. Interactions between the promoters and the genome in 15 kbp bins was counted with the connectCounts function. Interchromosomal interactions, interactions in blacklisted regions, with an anchor larger than 20 kbp or in the diagonal were excluded. A loess curve (span 0.1) was fitted to the average log counts per million (logCPM) for all libraries as a function of distance^0.25 (loessFit function from the limma package v3.40.2 (Ritchie et al., 2015)). An interaction was required to have an abundance larger than the fitted curved plus two times the mean of the absolute values of the residuals from the loess fit. Interactions for each promoter were then aggregated to produce a count matrix. Low-abundance promoters were filtered using edgeR’s filterByExpr function. Only promoters for protein-coding genes were retained. Obsolete Entrez Gene Ids were removed, as were mitochondrial, Riken, olfactory genes. The counts were normalized between libraries using a loess-based approach with the normOffsets function from the csaw package v1.18.0 (***Supp Fig S3***).

Differentially interacting promoters were detected with the quasi-likelihood (QL) framework (Chen et al., 2016; Lund et al., 2012) of the edgeR package. A design matrix was constructed with a one-way layout that specified the cell type. Using the promoter counts, the estimateDisp function was used to maximise the negative binomial likelihood to estimate the common dispersion across all promoters with trend=none and robust=TRUE (Chen et al., 2014). A generalized linear model (GLM) was fitted to the counts (McCarthy et al., 2012) and the QL dispersion was estimated from the GLM deviance with the glmQLFit function with robust=TRUE and trend.method=none. The QL dispersions were then squeezed toward a second mean-dependent trend, with a robust empirical Bayes strategy (Phipson et al., 2016) to share information between genes. A *P-*value was calculated for each promoter using a moderated t-test with glmQLFTest. The Benjamini– Hochberg method was used to control the false discovery rate (FDR) below 5%. Heatmaps of the filtered and normalized logCPM value were plotted with the coolmap function from the limma package. Gene set enrichment was tested using limma’s fry function and visualised using limma’s barcode enrichment plot.

### Overlaps between genomic features

Overlaps between genomic regions were identified with the overlapsAny function of the IRanges package v2.20.2 (Lawrence et al., 2013). Distances between nearest genomic features was calculated with the distanceToNearest function from the GenomicRanges package (Lawrence et al., 2013). Publicly available H3K27ac and H3K9me3 ChIP-seq, and DNaseI-seq data fastq files from wildtype DP thymocytes was downloaded from GSE63732. CTCF ChIP-seq and input data fastq files and from DP thymocytes was downloaded from GSE41743. Files were aligned with Rsubread with the mm10 reference genome. Duplicate reads were removed with Picard tools. Coverage plots and bedGraphs were created with Deeptools. The HOMER pipeline was used to call peaks (Heinz, et al 2010). Tag directories were created for each library with the genome specified. For the H3K27ac and H3K9me3 ChIP-seq, the findPeaks function was used to find peaks with style as histone and size as auto.

ChIP-seq and DNaseI-seq read coverage (log2 of the read per kilobase per million) of different genomic features was determined with featureCounts from Rsubread and the rpkm function from R package edgeR with log=TRUE. Read coverage of genes was determined from regions that are 5kbp upstream and 3kbp downstream from the TSS. Non differential expressed genes survived filtering in the DE analysis and adjusted p-value >0.05. For LADs, DIs and differential TAD boundary regions the genome regions excluded blacklisted regions. The p-value between groups was determined with a Mann-Whitney test.

### Motif enrichment

Motif enrichment was performed with the HOMER pipeline, on genomic regions with the findMotifsGenome.pl function and specifying the genome as mm10 and size given and for genes with the findMotifs.pl function with mouse specified as the genome. For DIs, enrichment was performed on the unique anchor regions of unclustered DIs relative to a background of the same number of not significant DI anchors (FDR>0.05).

### Identification of TAD cliques and modelling with Chrom3D

The Chrom3D pipeline was used to analyse the clique structure and model the 3D chromosome structure of the WT and DKO cells (Paulsen et al, 2017,2018) with some modifications. For creating the input GTrack file we used: intrachromosomal interactions at 100 kbp, interchromosomal interactions at 1 Mbp, the LADs as determined by EDD in the WT and DKO libraries separately and a diploid cell. Beads were created from TADbit segmentation at 50 kbp of summed biological replicates of the WT libraries excluding the Y chromosome with the boundary rounded down to the nearest 50 kbp. As there were no overlapping TADs, we did not merge beads. For calculating the significance of the intrachromosomal interactions we used fdr_bh with cutoff 1.5 and threshold 0.05 and for interchromosomal interactions cutoff 2 and threshold 0.05. TAD cliques were identified as previously described (Paulsen et al., 2019a). For each sample type we simulated one hundred independent Chrom3D models each with a separate seed value. We ran each simulation for 5 million iterations with a 5 μm radius, the nucleus flag and 0.3 scale. Bead measurements were calculated as the median radius of beads of each feature type for a given model. Distributions of these measurements were compared by Welch’s unequal variances t-test.

## Results

### Loss of heterochromatin results in predominant gene repression

In order to investigate the molecular consequences of heterochromatin loss, we performed RNA-seq on Suv39DKO and littermate control CD4^+^ CD8^+^ double positive (DP) thymocytes (***Supp Fig S1***), an abundant primary immune cell type that we have previously shown to be over 95% reduced in global H3K9me3 levels in the absence of both Suv39h enzymes (Keenan et al., 2020). Surprisingly, we identified a predominant down-regulation of gene expression with 78% of differentially expressed (DE) genes showing a negative logFC (***Fig 1a, Supp Table 2***). As expected both *Suv39h1* and *Suv39h2* were included in these DEs (***Supp Fig S4, Supp Table 2***).

**Figure 1.**
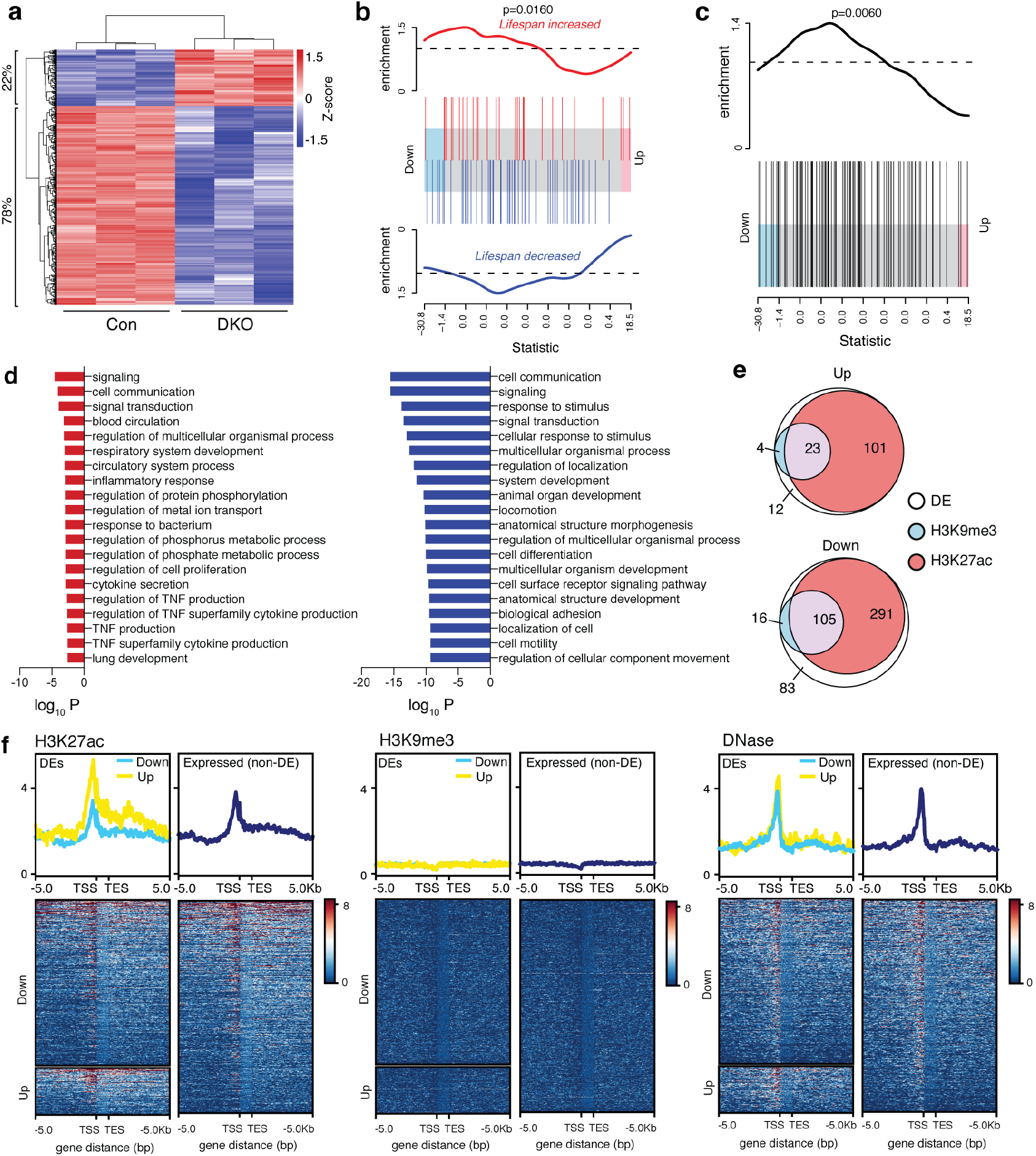
Heterochromatin loss causes predominant gene repression in euchromatic regions. **a)** Heatmap showing change of expression (logRPKM) of differentially expressed (DE) genes between Suv39h1 and h2 double knockout (DKO) and control cells. The proportion of genes down-regulated and up-regulated in Suv39DKO cells versus control are annotated. **b)** Barcode enrichment plot showing ranking of ageing-related genes from the GenAge database (de Magalhaes et al., 2009) amongst the DE genes. Genes are ranked right to left from most up-regulated to most down-regulated in DKO cells. The rank of genes associated with increased lifespan is marked by read vertical bars and that of genes associated with decreased lifespan by blue vertical bars. Red and blue worms show relative enrichment. ROAST gene set test p-values tests correlation. **c)** same as (b) but with directionality removed to include genes with contradictory annotated life-span effects. **d)** gene ontology (GO) enrichment in DE genes. GO terms enriched in up-regulated genes shown in red and down-regulated genes shown in blue. **e)** quantification of overlap between DE genes and H3K27ac and H3K9me3 ChIP-seq peaks as called by HOMER. **f)** coverage plots of DE and non-DE genes showing H3K27ac, H3K9me3 and DNaseI accessibility levels from 5 kb upstream of the transcription start site (TSS) to 5 kb downstream of the transcription end site (TES).

Given heterochromatin loss has been linked to premature ageing in studies by us (Keenan et al., 2020), and others (Larson et al., 2012; Zhang et al., 2015)., we performed gene-set enrichment analysis on ageing-related genes from the GenAge Database (de Magalhaes et al., 2009). We noted many ageing-related genes have duplicated entries with conflicting effect annotations (i.e. annotated to both increase and decrease lifespan). Using only genes with unique annotation, we found a positive correlation (P=0.0165) between the ageing signature and the DE genes suggesting genes that increase lifespan tend to be up-regulated in the DKO, while those that decrease lifespan tend to be down-regulated. However, examination of the resultant barcode plot (***Fig 1b***) reveals this correlation is predominantly driven by down-regulated genes. When we include all ageing-related genes (to include those genes with contradictory annotated life-span effects) we find a similar enrichment in down-regulated genes (***Fig 1c***).

In order to investigate whether the gene regulation observed in the Suv39DKO was a result of secondary effects such as through the dysregulated expression of particular transcription factors (TF), we performed TF binding motif analysis. However, we found no significantly enriched motifs in either our up- or down-regulated gene sets. We next performed gene ontology analysis revealing many statistically enriched pathways for both up-regulated and down-regulated genes (***Fig 1d***). We noted that these pathways appear to be general cellular processes, rather than ageing-related, chromatin-related or immune-system specific processes suggesting that the gene regulation is not a result of disruption of a particular cellular pathway.

### Genes regulated following heterochromatin loss lie in euchromatic regions, despite a loss of lamina-associated domains (LADs)

We were intrigued by the predominant down-regulation of genes following heterochromatin loss so we next used publicly available ChIP-sequencing and DNaseI-sequencing data from wildtype DP thymocytes (Vanhille et al., 2015) to see whether the DE genes lie in open (DNaseI accessible), active (H3K27ac-marked) or repressive (H3K9me3-marked) regions. Remarkably, the DE genes following heterochromatin loss predominantly lie in DNaseI accessible, H3K27ac-marked regions, rather than repressive H3K9me3-marked regions (***Fig 1e,f***), suggesting heterochromatin loss is affecting euchromatic gene expression.

Given the role of the Suv39h-H3K9me3-HP1 pathway in facilitating association of heterochromatic domains with the nuclear periphery, we next performed LaminB1 ChIP-seq on control and Suv39DKO DP thymocytes. Using the Enriched domain detector algorithm (Lund et al., 2014) specifically developed for Lamin ChIP-seq to call LAD domains, we found a reduced number of LADs in DKO cells, and those LADs which were retained were significantly smaller in size compared to control cells (***Fig 2a-c, Supp Table 3***). In accordance with our observations that DE genes predominantly fall in euchromatic regions (***Fig 1e,f***), only a small minority of DE genes overlapped LaminB1-associated regions for either up- or down-regulated genes (***Fig 2a,d).*** Given the canonical role of heterochromatin in gene silencing, we were surprised to find that a greater proportion of down-regulated genes in LAD-disrupted regions compared to upregulated genes in terms of both number of DE genes (***Fig 2d***) and logFC (***Fig2e***). As expected, LADs (both lost and retained in the DKO) exhibited very little H3K27ac levels whereas LADs disrupted in the DKO were marked by higher levels of H3K9me3 compared to LADs which were retained in the DKO, or the rest of the genome (***Fig 2f***).

**Figure 2.**
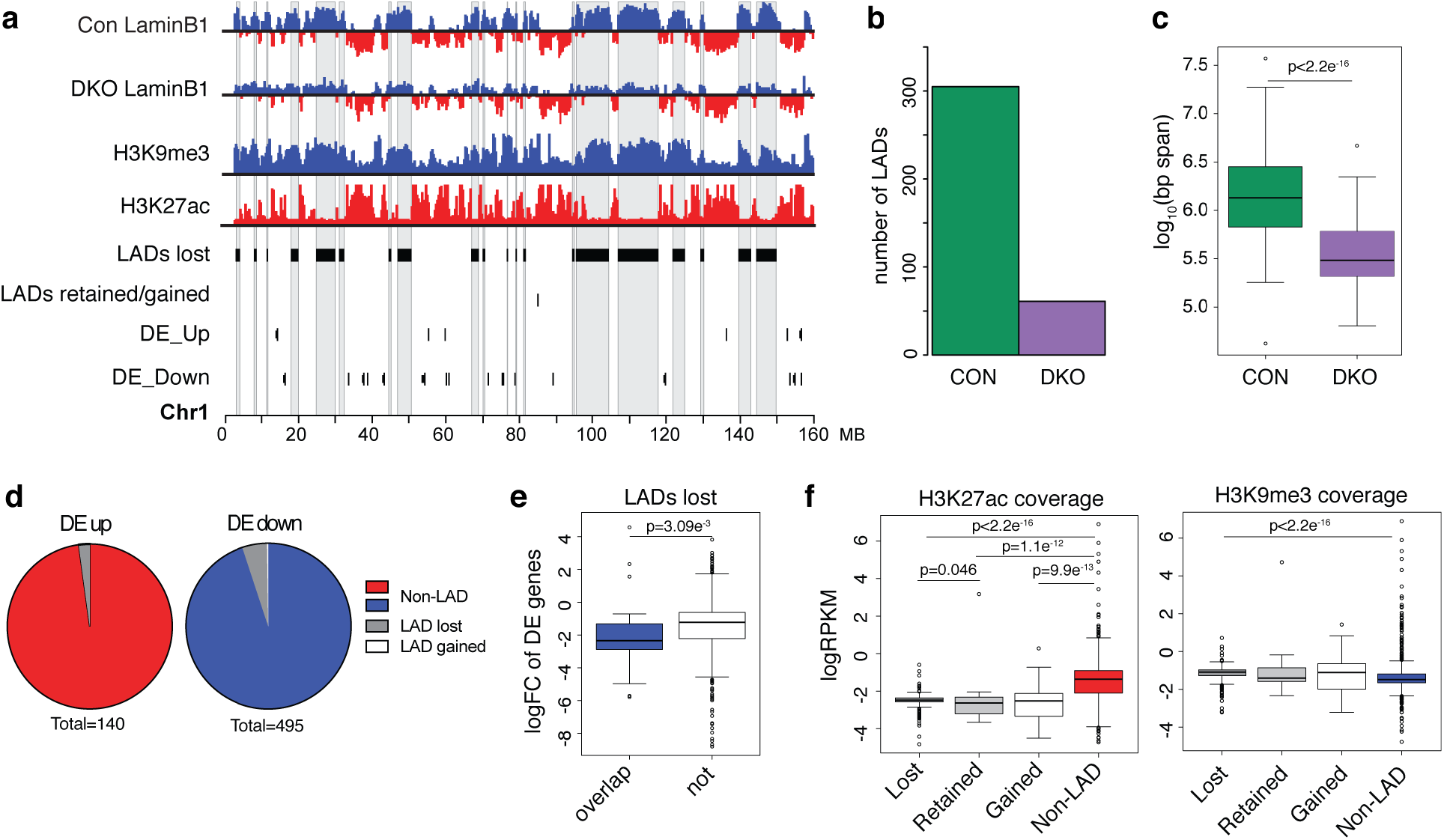
Suv39DKO reduces lamina-associated domains (LADs) without inducing gene activation. **a)** Integrative Genomics Viewer tracks of LaminB1 ChIP-seq from control and Suv39DKO DP thymocytes overlayed with H3K9me3 and H3K27ac ChIP-seq from wildtype DP thymocytes (Vanhille et al., 2015) and DE genes. Lamina-associated domains (LADs) are divided into LADs lost in DKO and LADs retained or gained. Chromosome 1 (0 - 160 MB) is shown with ChIP-seq tracks at 100 kb resolution. **b)** Number of LADs called across all chromosomes in control and Suv39DKO DP thymocytes. **c)** Distributions of LAD size in control and Suv39DKO DP thymocytes. Box plot depicts the interquartile range (IQR) ± 1.5xIQR and median annotated. Distributions were compared by Wilcoxon rank sum test with continuity correction. **d)** proportion of DE genes between Suv39DKO and control cells which overlap differential LADs. **e)** Distribution of logFC for DE genes overlapping LADs lost in Suv39DKO cells compared to control. Distributions shown as box plots as in (**c)** and statistically compared by Wilcoxon rank sum test with continuity correction. **f)** Distributions of H3K27ac and H3K9me3 logRPKM across LADs lost, gained and retained in the Suv39DKO versus the rest of the genome. Distributions shown as box plots as in (**c)** and statistically compared by Wilcoxon rank sum test with continuity correction.

### Heterochromatin loss causes switching of active to repressive compartments

To investigate how other forms of genome organization other than lamina-association were altered following the loss of heterochromatin, we performed *in situ* Hi–C on Suv39DKO and control DP thymocytes. We obtained high quality libraries of approximately ~120 million read pairs per biological replicate (after filtering) with two independent replicates per condition (***Supp Table 1***). First, we examined the large scale genomic compartments of the interaction space comprising euchromatic, gene dense, so-called A compartment and the gene poor, heterochromatic, B compartment. We found a relatively large amount of compartment switching between the Suv39DKO and control DP thymocytes (~1.5% of the genome) (***Fig 3a***). However, given the loss of lamina-associated domains in the Suv39DKO (***Fig 2a-c***), we were surprised to find that most of this compartment switching was from an active A compartment in control cells to a repressive B compartment in the DKO (***Fig 3b***). We found that no up-regulated DE genes fall in compartment switched regions (***Fig 3c***). However, a subset of down-regulated genes (7.5%) overlap compartment switched regions, with all of these regions switching from A to B (***Fig 3c***) consistent with the decrease in expression of these genes.

**Figure 3.**
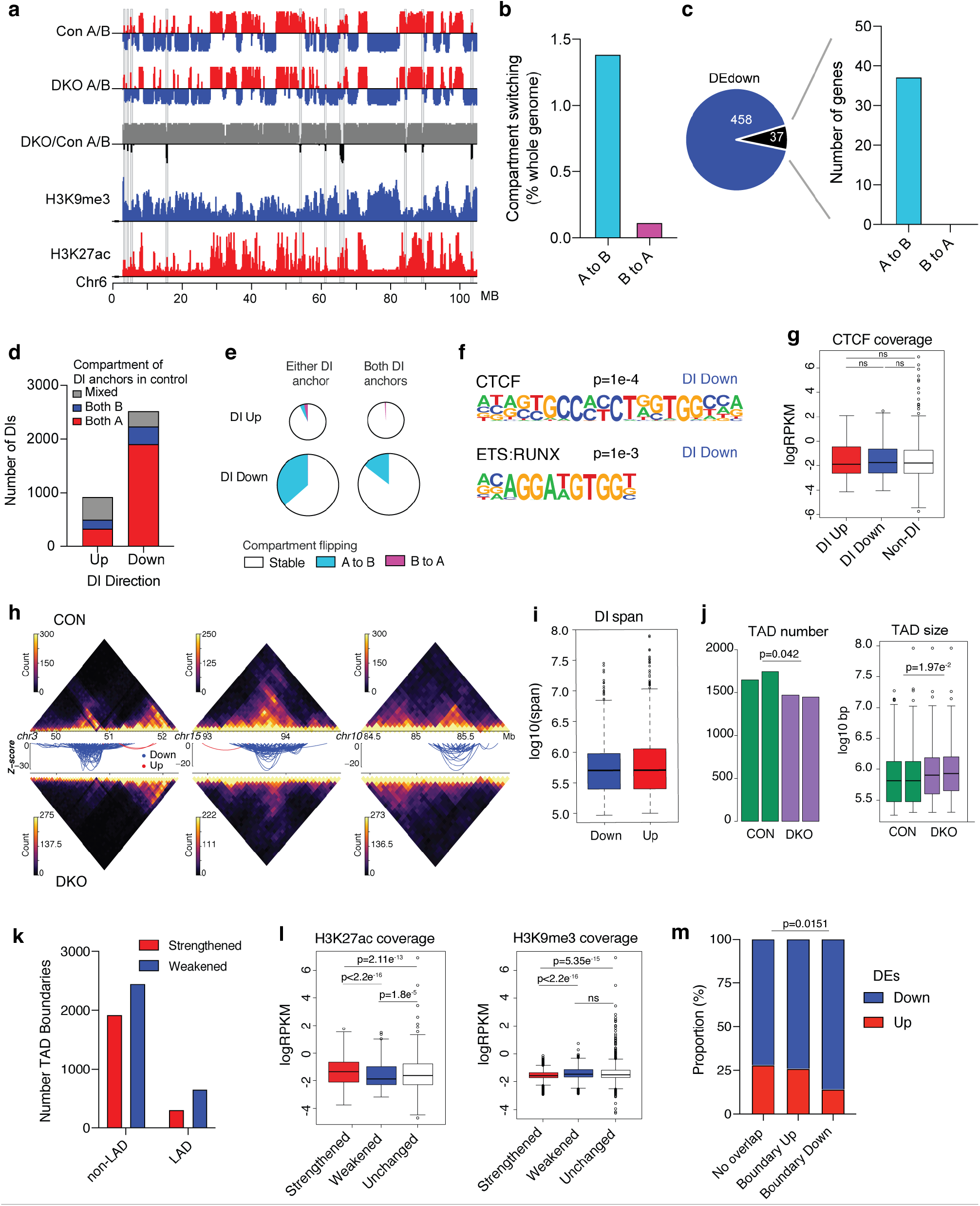
Heterochromatin loss causes a loss of chromatin interactivity in active regions and significant switching from active to repressive compartments. **a)** Integrative Genomics Viewer tracks of A/B compartments from Hi-C analysis of Control and Suv39DKO DP thymocytes overlayed with H3K9me3 and H3K27ac ChIP-seq from wildtype DP thymocytes (Vanhille et al., 2015). Chromosome 6 (0 – 105 MB) is shown at 100 kb resolution. **b)** proportion of genome that switches A/B compartments in Suv39DKO versus control cells (shown as % of whole genome). **c)** overlap of down-regulated DE genes with compartment switched regions. **d)** number of unclustered differential interactions (DIs) (FDR<0.05) between Suv39DKO and control cells. Strengthened interactions (logFC>0) are annotated as ‘up’ and weakened (logFC<0) as ‘down’. Overlap of the DI anchors with compartment A and B in control cells in shown. DIs where both anchors are not contained in the same compartment, or where one or more anchor overlaps both compartments, are annotated as ‘mixed’. **e)** The proportion of DIs with anchors which overlap switched compartments. **f)** transcription factor binding motifs enriched in the anchors of DIs as determined by the HOMER pipeline. **g)** CTCF ChIP-seq (Shih et al., 2012) coverage of the anchors of DIs versus the rest of the genome. Box plot depicts the interquartile range (IQR) ± 1.5xIQR and median annotated. Distributions were compared by Wilcoxon rank sum test with continuity correction. **h)** Hi-C contact matrices at 50kbp resolution showing the top 3 DI regions between Suv39DKO and control cells. Colour scale indicates the number of read bins per bin pair with visualisation scaled to total library size to allow appropriate visual comparison. Unclustered DIs (FDR <0.05) are shown as arcs (blue indicate a decrease in logFC, red an increase in logFC) where the vertical axis is the −log10(p-value) of the DI. **i)** the linear span between genomic anchors of strengthened (‘up’) and weakened (‘down’) DIs. Data shown as boxplot as in (**g**). **j)** number and size of topologically associated domains (TADs) in each replicate sample of control and Suv39DKO cells. Data statistically compared by unpaired t-test with equal variance between the median of the TAD sizes and number of TADs. Boxplot for TAD size plotted as in **(g)**. **k)** Number of TAD boundary changes between Suv39DKO and control cells divided into those overlapping LAD and non-LAD regions in control cells. **l)** H3K27ac and H3K9me3 density (shown as logRPKM) of strengthened and weakened TAD boundaries in Suv39DKO and control cells versus the rest of the genome. Data shown as boxplot as in (**g**) and compared by Wilcoxon rank sum test with continuity correction. **m)** proportion of DE genes up-regulated versus down-regulated overlapping altered TAD boundaries and the rest of the genome. Data analysed by Chi-squared test.

### Heterochromatin loss weakens genome organization in euchromatic regions, particularly in TAD boundaries

We next utilized the *diffHic* package (Lun and Smyth, 2015b) to identify discrete regions of altered 3-dimensional genome organisation between Suv39DKO and control cells. *diffHic* uses the statistical framework of the edgeR package (Robinson et al., 2010) to model biological variability and test for significant differences (termed differential interactions, DIs). Using a resolution of 50 kb, we found >3000 DIs (FDR<0.05) between Suv39DKO and control cells with the majority of DIs identified as a weakening of structure (logFC<0) rather than a gain of structure (***Fig 3d, Supp Table 4***). We found that the majority of weakened DIs had both anchors (the physically interacting genomic regions mapped from the Hi-C read pairs) of the DI in the A compartment (***Fig 3d***). Furthermore, a significant proportion of weakened DIs overlapped A to B compartment switched regions even when only one DI anchor was in this region (***Fig 3e***).

We next performed TF motif analysis in the anchors of the DI loops in order to identify putative regulators mediating these alterations in 3-dimensional genome organisation. We found 2 motifs to be significantly enriched in weakened DI anchors including that of CCCTC-binding factor (CTCF), the canonical regulator (together with cohesin complex) of chromatin architecture (Ong and Corces, 2014; Rao et al., 2014; Tang et al., 2015) (***Fig 3f***). To explore whether CTCF occupancy affects which regions alter structure in the Suv39DKO we overlayed downloaded publicly available CTCF ChIP-seq from DP thymocytes (Shih et al., 2012). However, we found no evidence that CTCF binding was enriched or depleted in structure altered in the Suv39DKO (***Fig 3g***).

Visualisation of our top differential interacting regions revealed that the altered structure in the Suv39DKO are of an impressive scale in the order of entire topologically associated domains (TADs) at 0.4 - 1 Mb size (***Fig 3h,i***). We therefore next explored whether TAD structures are altered in the Suv39DKO using the TADbit package (Serra et al., 2017). We found slightly reduced TAD numbers in the DKO and a commensurate increase in average TAD size (***Fig 3j***). To further explore the dynamics of TAD changes we identified regions of changing contrast, i.e., strengthening or weakening of TAD boundaries. The TAD caller TADbit uses a binary approach that cannot identify subtle changes in chromatin architecture such as the weakening of a boundary as opposed to its complete absence. Therefore, we employed a different approach to identify regions where the TAD boundaries change in strength by adapting the statistical strategy recently described for differential methylation (Chen et al., 2017) and the *diffHic* package (Lun and Smyth, 2015a). This method directly assesses differential boundary strength relative to biological variation without needing to make TADs calls in individual samples. We first segmented the entire genome into 200 kb regions and then assessed whether the boundary strength at this region changed by assessing the contrast in up-stream and down-stream intensity. The ratio of up-stream to down-stream intensity in Suv39DKO and control cells was then compared using edgeR in a statistically robust manner.

We found that TAD boundaries are more frequently weakened in the Suv39DKO regardless of whether the TADs are lamina-associated or not (in control cells) (***Fig 3k, Supp Table 5***). Interestingly, those TAD boundaries that strengthened in the DKO were marked by higher H3K27ac levels and lower H3K9me3 levels in control cells compared to boundaries which weakened or were unchanged, whereas TAD boundaries which were strengthened were marked by lower H3K27ac levels (***Fig 3l***). Critically, 62% of DE genes fall in genomic regions where these TAD boundaries are modulated. Exploring the directionality of differential gene expression in relation to the strengthening or weakening of TAD boundaries, we found that genes which overlap weakened boundaries are more likely to be repressed compared to DE genes which overlap strengthened TAD boundaries, or DE genes that fall in other genomic loci (***Fig 3m***).

### Higher-order modelling of the 3D nucleome in Suv39DKO cells reveals heterochromatin loss affects the expression of genes positioned closer to the nuclear periphery in 3-dimensional space

We next explored whether higher-order chromatin organisation was altered in Suv39DKO cells by assessing the clustering of non-contiguous TADs into so-called cliques (Paulsen et al., 2019b). As has previously been reported in adipose progenitor cells, we could detect TAD cliques containing up to 11 interacting TADs in control DP thymocytes (***Fig 4a***). However, in Suv39DKO cells the detected TAD cliques near-uniformly contained fewer interacting TADs (***Fig 4a,b***). Interestingly, in our data from DP thymocytes, the majority of TAD cliques were found in compartment A (***Fig 4a,c***) in contrast to what has previously been observed in adipose cells (Paulsen et al., 2019b). In the Suv39DKO we saw a similar numerical reduction in the number of cliques from both compartment A and B (***Fig 4c***). However, given cliques are predominantly in compartment A in control cells, those in compartment B were proportionally much more disrupted in the Suv39DKO (***Fig 4c***).

**Figure 4.**
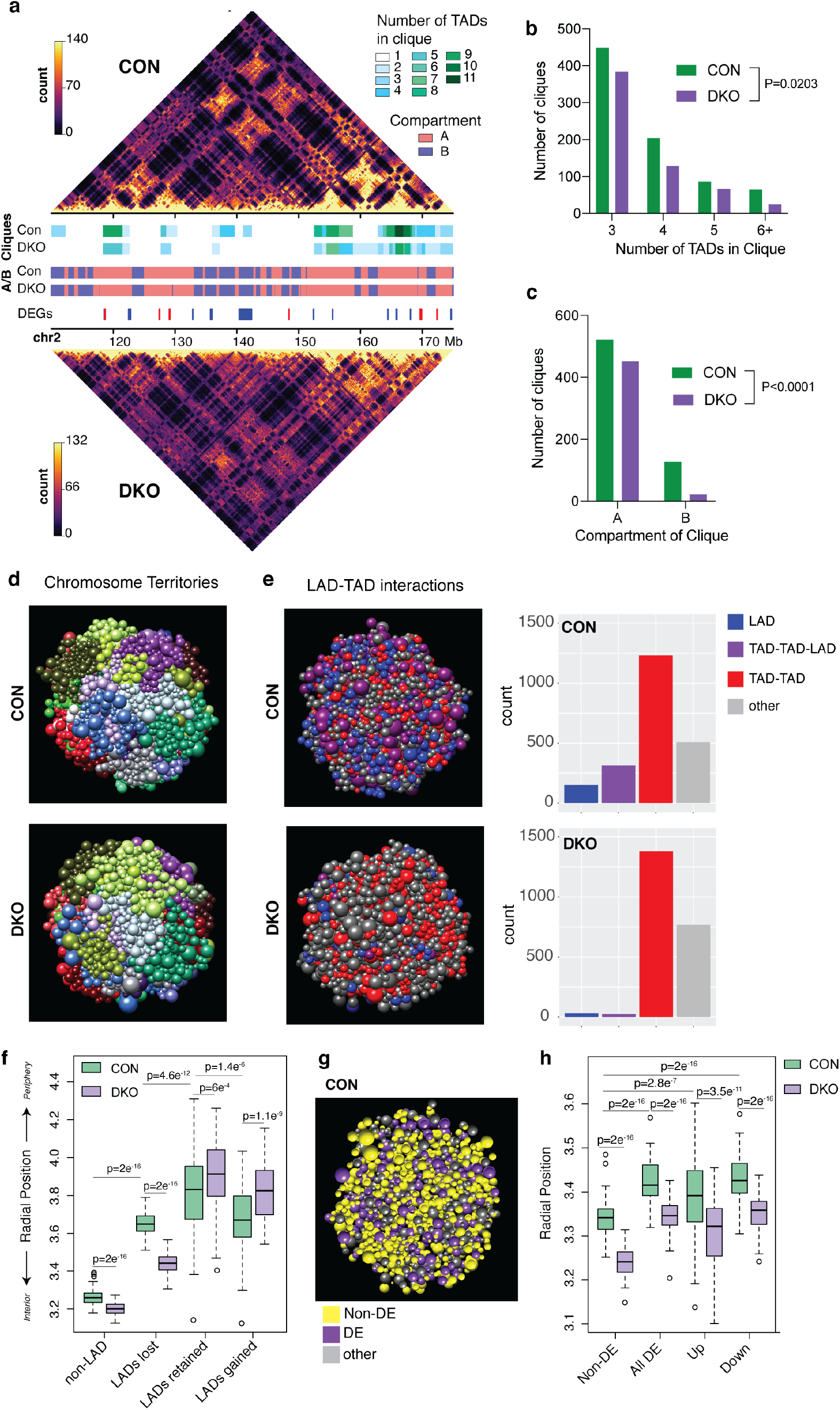
Higher-order modelling of the 3D nucleome in Suv39DKO cells reveals heterochromatin loss affects the expression of genes positioned closer to the nuclear periphery in 3-dimensional space. **a)** Representative Hi-C contact matrices at 200kbp resolution showing a loss of higher-order TAD-TAD cliques in Suv39DKO cells. Colour scale indicates the number of read bins per bin pair with visualisation scaled to total library size to allow appropriate visual comparison. A/B compartments are also annotated as are differentially expressed genes (DEGs, blue is downregulated and red is upregulated). **b)** Number of higher-order TAD-TAD cliques detected in Suv39DKO and control cells. Data statistically compared by 2-way ANOVA. **c)** Number of higher-order TAD-TAD cliques in compartment A and compartment B in Suv39DKO and control cells. Proportion of cliques in each compartment data statistically compared by Chi-squared test. **d,e)** Representative Chrom3D modelling of the 3D nucleome of control and Suv39DKO cells coloured as individual chromosome territories **(d)** or higher-order TAD-TAD, TAD-TAD-LAD or LAD interactions **(e)**. **f)** Measurements of LAD positioning from Chrom3D modelling in Suv39DKO and control cells. Box plot depicts the interquartile range (IQR) ± 1.5xIQR with median annotated from 50 independent modelling iterations each from a separate seed value. Distributions were compared by Welch’s unequal variances t-test. **g,h)** Measurements of DE gene positioning from Chrom3D modelling as in **(f)**. Distributions were compared by Welch’s unequal variances t-test.

We next modelled the 3D nucleome of control and Suv39DKO cells using the computational platform *Chrom3D* (Paulsen et al., 2018; Paulsen et al., 2017). Chrom3D simulates the spatial position of chromosome domains as defined by TADs relative to each other from the Hi-C data, and relative to the nuclear periphery from LaminB1 ChIP-seq data. We produced 50 independent models for both control and Suv39DKO cells (an example model for each cell type is shown in ***Fig 4d***). We examined higher-order structure of the chromosome domains including LADs, TAD-TAD interactions and TAD-TAD-LAD interactions revealing a loss of TAD-TAD interactions (demonstrated by an increase in ‘other’ counts in the Suv39DKO), as well as a loss of TAD-TAD interactions from the nuclear periphery (***Fig 4e***).

Quantification of the median position of domains with LAD interactions in each independent simulation of the control and Suv39DKO nucleome accord with our expectations showing LADs in control cells positioned closer to the nuclear periphery than non-LADs regardless of whether they are lost, retained or gained in the Suv39DKO (***Fig 4f***). Moreover, LADs lost in the Suv39DKO show a highly significant shift towards the nuclear interior (***Fig 4f***). Most interestingly, when we then overlayed our DE genes onto the *Chrom3D* models we found that DE genes were positioned closer to the nuclear periphery than expressed non-DE genes (***Fig 4g,h***), regardless of whether they were up- or down-regulated (***Fig 4h***).

### Altered 3-dimensional genome organisation in LAD proximal regions correlates with transcriptional regulation

We were intrigued by the mechanism by which these genes near the nuclear periphery were affected by the loss of heterochromatic structure. We hypothesised that the loss of LADs could be disrupting the 3-dimensional chromatin interactivity around nearby genes thereby disrupting their expression. We therefore adapted our recently developed gene-centric differential interaction analysis (Chan et al., 2019) to explore this hypothesis. This approach calculates interaction counts between the promoters of coding genes and the rest of the genome in 15 kbp bins, filters these interaction counts using a distance-dependent approach, then aggregates these counts for each gene. Gene-centric differential interactivity is then detected between samples using the quasi-likelihood (QL) framework of the edgeR package (***Fig 5a***). Interestingly, we found that chromatin interactivity showed a similar pattern to transcriptional regulation (***Fig 1a***), with interactivity at coding genes more frequently reduced in Suv39DKO cells (***Fig 5b, Supp Table 6***). Similarly, interactivity around ageing-associated genes showed a significant enrichment for weakened interactivity in this gene set (***Fig 5c***), consistent with what we observed for gene expression (***Fig 1c***). We next examined whether differential interactivity was related to proximity to the nuclear periphery. To do this, we divided the genome into bins based on the linear distance to the closest LAD (in control cells) and explored the differential interactivity (as LogFC between Suv39DKO and control) of genes within each bin. We found that genes on LADs show the largest loss of interactivity in Suv39DKO cells (although this gene number is relatively few), with proximal genes showing an intermediate loss of interactivity with this effect being lost around 3Mb from the nearest LAD (***Fig 5d***). Critically, we found a significant correlation between the altered expression of genes in Suv39DKO cells and the altered interactivity of these genes (***Fig 5e***) with interactivity predominantly lost and gene expression predominantly repressed (***Fig 5f***). This suggests that the loss of lamina-associated heterochromatin could indeed be disrupting euchromatic gene expression through a disruption of nearby 3-dimensional chromatin interactivity (***Fig 5g***).

**Figure 5.**
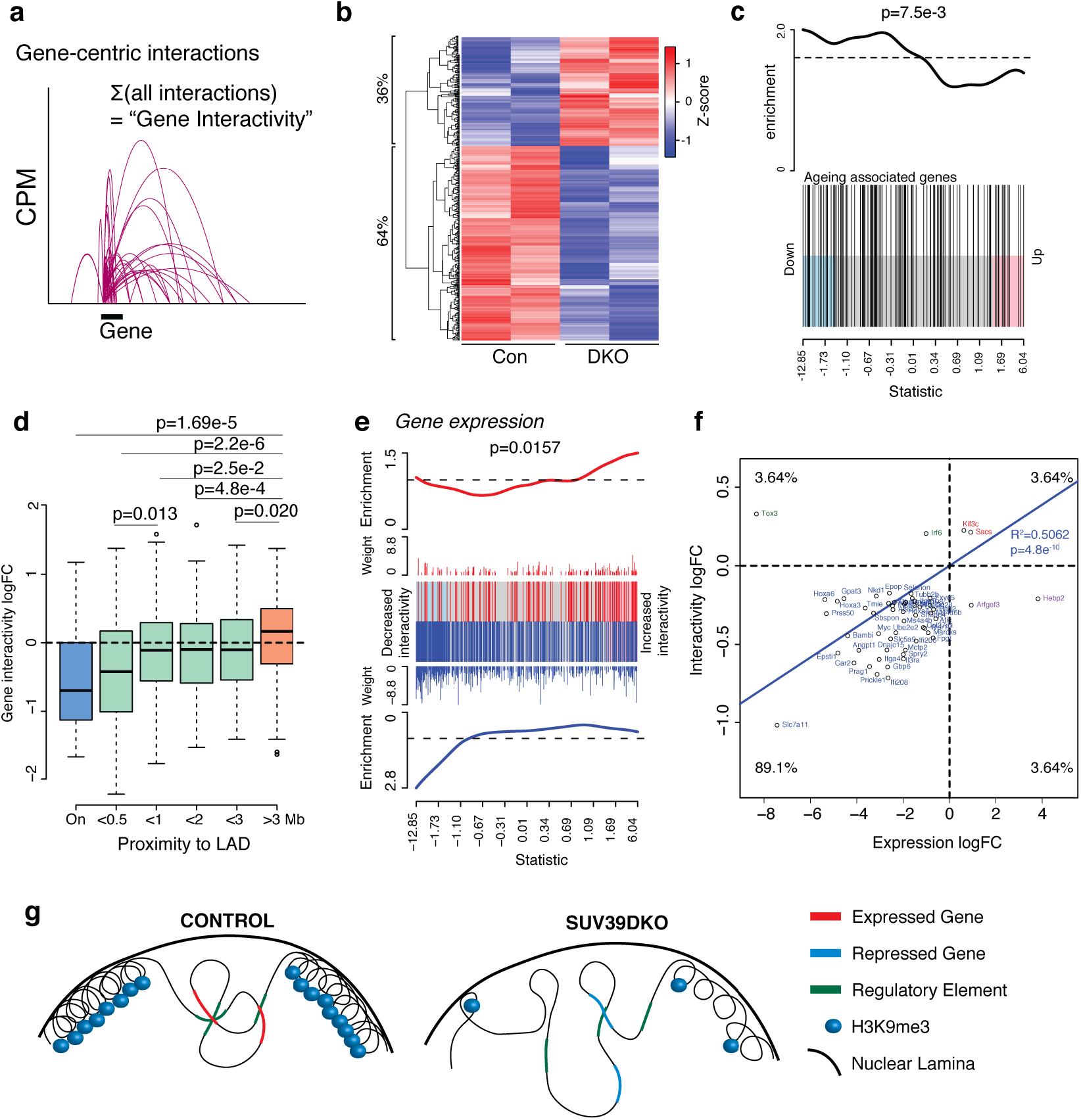
Altered euchromatic genome interactivity near LAD domains correlates with transcriptional dysregulation in Suv39DKO cells. **a)** Schematic of gene-centric interactivity analysis. **b)** heatmap of the interactivity (log2CPM) of differentially interacting genes between Suv39DKO and control cells (FDR<0.05). **c)** Gene Set Enrichment Analysis of genes with differential interactivity relative to ageing-related genes from the GenAge database. Barcode enrichment plot shows the correlation of ageing-related genes relative to differential interactivity. Genes are ordered on the plot from right to left (x-axis) from most increased in interactivity to most decreased in interactivity according to the moderated F-statistic. P-value was calculated with the fry test. **d)** logFC of gene-centric interactivity for DE genes shown relative to linear genomic distance to the nearest lamina-associated domain (LAD) and statistically compared by Wilcoxon rank sum test with continuity correction. **e)** barcode enrichment plot showing the correlation of differentially expressed genes relative to differential interactivity. Genes are ordered on the plot from right to left (x-axis) from most upregulated to most downregulated according to the moderated F-statistic. P-value was calculated with the fry test. **f)** logFC of gene-centric interactivity relative to logFC of transcriptional change for DE genes. **g)** schematic of putative mechanism by which the loss of heterochromatin causes transcriptional repression.

## Discussion

We have here revealed that loss of Suv39-dependent heterochromatin domains causes predominant gene repression, with differentially expressed genes principally in euchromatic regions rather than heterochromatic regions. We found that differentially expressed genes are closer to the nuclear periphery and highly correlated with altered 3-dimensional genome interactivity which we believe is driven by the loss of lamina-association of Suv39DKO chromatin. Together, our results suggest that the nuclear lamina-tethering of Suv39-dependent H3K9me3 domains provides an essential scaffold to support euchromatic genome organisation and the maintenance of gene transcription for healthy cellular function.

It is now well accepted that most genes within a genome are controlled by long-range *cis*-regulatory elements physically looping to interact with their target gene, even though these regulatory elements may lie many kilobases or megabases away (Dekker et al., 2013; Dixon et al., 2015; Sanyal et al., 2012). Our data showed a high level of concordance between differential gene expression and differential interactivity of that gene suggesting either that gene expression *per se* is highly correlated to the interactivity of that gene with other genomic elements such as enhancers or other gene promoters, or that the mechanism causing gene expression to alter similarly affects the interactivity around that gene. Given there are many complex regulatory mechanisms controlling the expression of individual genes, it seems implausible that chromatin interactivity could be the sole determinant of gene expression. Indeed, a recent study exploring *cis*-regulatory elements through ATAC-seq profiling has found that whilst the expression of most genes are dependent on long-range enhancers, a significant fraction are not, particularly for housekeeping and cell-cycle related genes (Yoshida et al., 2019). Moreover, a study exploring chromatin interactivity and gene expression in T cell development found that only a subset of genes showed correlation between interactivity and expression, although these genes were deemed critical to the T cell development process (Hu et al., 2018). We therefore contend that the mechanism by which the loss of Suv39-dependent heterochromatin causes altered transcription similarly affects the interactivity around that gene.

The temporal or causal relationship between chromatin interactivity and gene transcription has been challenging to untangle due to the complex interdependency between the two processes. Some groups propose that genome reorganisation often precedes gene expression changes (Stadhouders et al., 2018), whereas others propose that these interactions form concomitant with gene expression (Oudelaar et al., 2020). Here, we propose that the structural alterations in chromatin organisation caused by a loss of LADs in the Suv39DKO are the driving force for the transcriptional change, and are not a consequence of transcriptional alterations. Whilst we have not demonstrated direct causality, we provide evidence that the magnitude of alteration of genome interactivity is inversely proportional to the proximity to a LAD and we believe that this is readily explained by biophysical connectedness. It may be argued that we also show differentially expressed genes to be similarly proximal to the nuclear periphery through our modelling of the 3D nucleome. However, it is less obvious what mechanism would affect the transcription of nearby genes in the absence of an intermediate effect on genome organisation. Increased levels of DNA damage may be a contributing factor (Nava et al., 2020). Indeed, we have previously shown that γH2AX levels are increased in bone marrow cells from Suv39DKO chimeric mice. However, it is not clear that DNA damage would result in the predominant down-regulation of gene expression we have observed in the absence of an intermediate effect on genome organisation. We explored the hypothesis that dysregulation of particular transcription factors may cause the altered gene transcriptional profile in the Suv39DKO given our lab and others have shown that TFs can alter both gene transcription and genome organisation (Hu et al., 2018; Johanson et al., 2018; Stadhouders et al., 2018). However, we found no enriched transcription factor motifs in the promoters of the differentially expressed genes. Therefore we believe that the dysregulated gene expression we observe is not a result of secondary effects from transcription factor dysregulation.

Our findings here using our unique model of heterochromatin loss in primary immune cells complement several recent papers in different models and systems. In particular, a recent study used mechanical stress to disrupt heterochromatin stability in skin epidermis stem/progenitor cells (Nava et al., 2020). They also found a predominant down-regulation of gene expression, reduced lamina-association of heterochromatin domains, but were unable to find a correlation between H3K9me3-regulated regions and gene transcriptional changes (Nava et al., 2020). A separate study examined the formation of senescence-associated heterochromatin foci (SAHF) in the oncogene-induced senescence of fibroblast cells where they found a gain of interactivity within 3MB of a newly formed heterochromatin domain (Sati et al., 2020). This perfectly complements our findings that genes within 3MB of LAD showed an overall decrease in interactivity compared to genes further away. Moreover, our conclusions are broadly complementary to the recent study suggesting heterochromatic regions drive the phase separation of active and inactive chromatin (Falk et al., 2019).

We have here revealed that down-regulated genes in the Suv39DKO are enriched for ageing-associated roles providing a new perspective on the potential molecular causes of cellular dysfunction in ageing. This is not to suggest that a predominant down-regulation of genes occurs in physiological ageing as we have seen here, as many studies have shown that different cell types and organisms do not show such a clear transcriptional effect but rather show an increased transcriptional variability (Bahar et al., 2006; Martinez-Jimenez et al., 2017). We instead contend that the molecular consequences of heterochromatin loss are more complex than a simple derepression of silenced elements, and that this transcriptional regulation in LAD-adjacent regions occurs with ‘physiological’ heterochromatin instability as is seen in ageing, it was just hitherto unseen. Given we have focused on the transcriptional regulation of protein-coding genes we do not know whether repetitive elements such as transposable elements have been activated as has been observed in other contexts (De Cecco et al., 2013a; De Cecco et al., 2013b; Li et al., 2013; Wood et al., 2016). However, it is likely that multiple molecular mechanisms act in concert to result in the cellular dysfunction associated with ageing. Through adding to our understanding of the molecular consequences of heterochromatin loss we hope intervention strategies may be designed to restore cellular function and prevent age-associated diseases.

## Acknowledgements

We thank Thomas Jenuwein (MPI, Frieburg) for the Suv39h-deficient mice (Peters et al., 2001). We acknowledge the tremendous technical assistance from the staff of the core facilities at the Walter and Eliza Hall Institute, particularly Stephen Wilcox in the WEHI genomics hub, the staff of the Flow Cytometry facility (WEHIFACS) and WEHI Bioservices. This work was supported by grants and fellowships from the National Health and Medical Research Council of Australia (CRK #1125436, TMJ #1124081, RSA #1100451), and the Australian Research Council (RSA #130100541). This study was made possible through Victorian State Government Operational Infrastructure Support and Australian Government NHMRC Independent Research Institute Infrastructure Support scheme.

## Author contributions

C.R.K., N.I., T.M.J., W.F.C. performed experiments; H.D.C., A.L.G performed formal analysis; C.R.K., H.D.C., R.S.A wrote the paper; G.K.S., R.S.A. supervised the research.

## SUPPLEMENTARY FIGURES

**Supplementary Figure S1:**
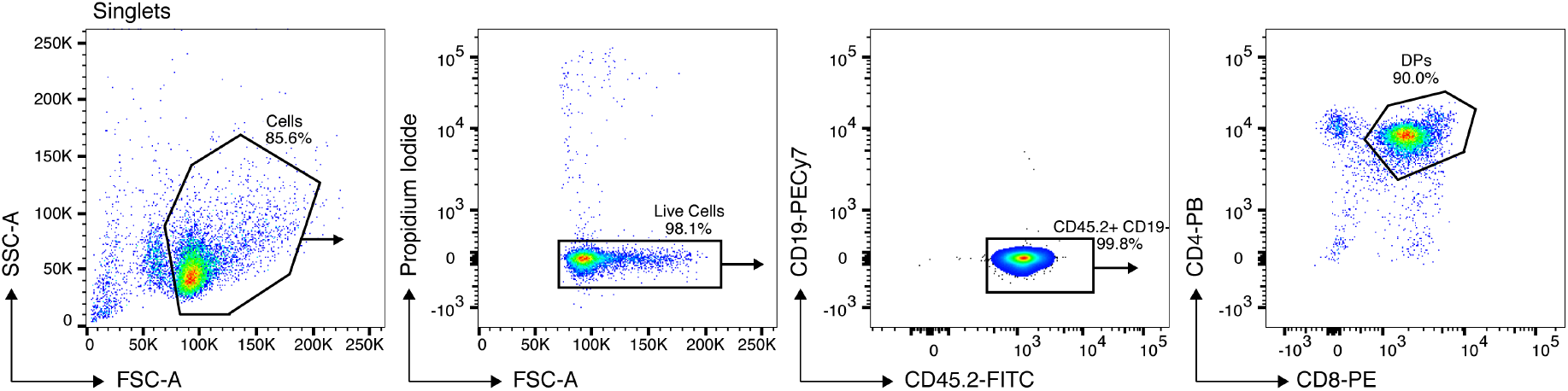
Representative flow cytometric gating of double-positive (DP) thymocytes. Primary DP thymocytes (CD4^+^CD8^+^CD19^−^) were flow-sorted from total thymic cells from Suv39DKO and control chimeric mice. The CD45.2^+^ congenic mark was used to identify donor cells of appropriate genotype as opposed to residual CD45.1^+^ white blood cells from the irradiated host.

**Supplementary Figure S2:**
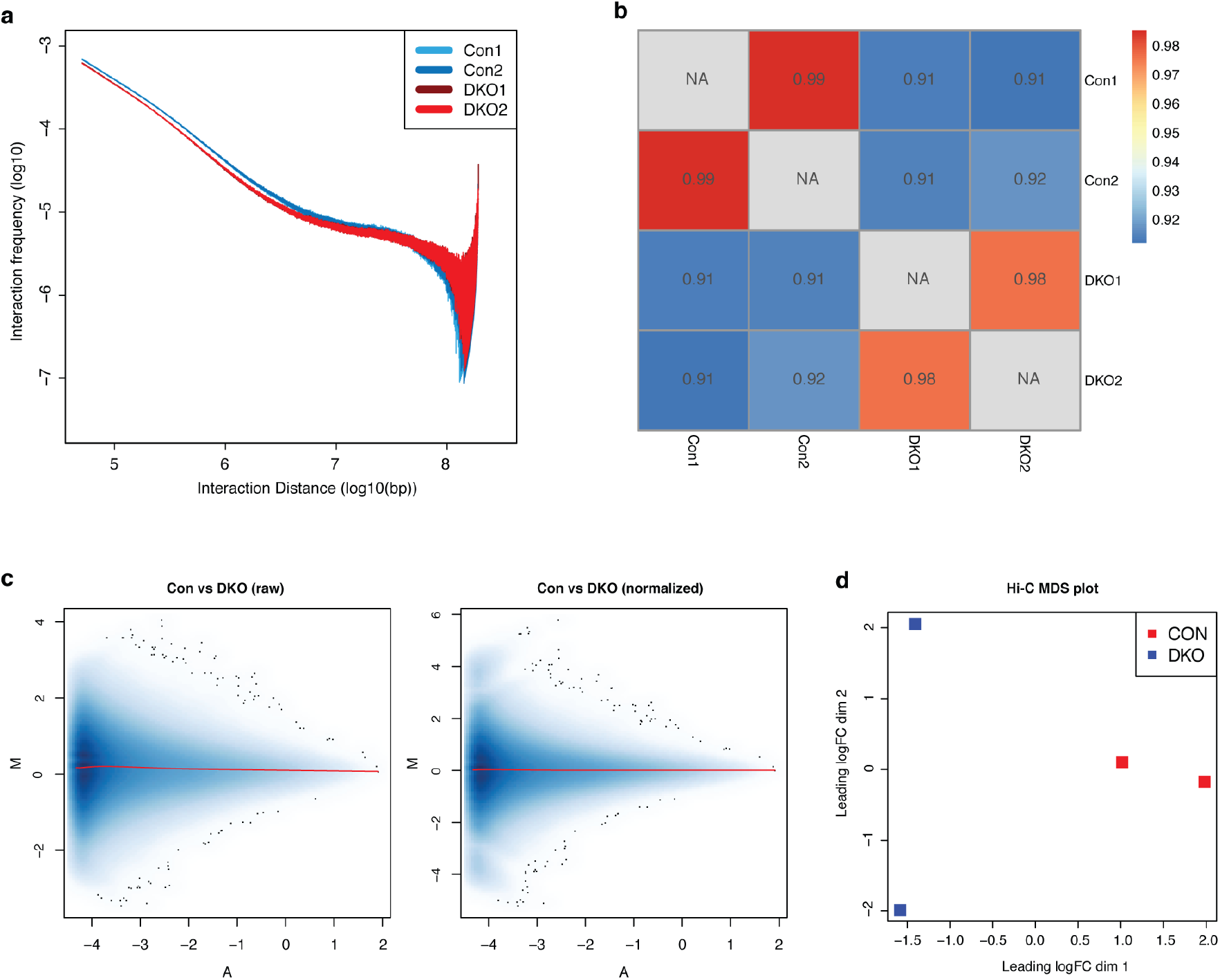
Normalization and reproducibility of HiC data (related to Figure 3). **a)** Interaction decay curves of libraries of the read-pair interaction frequency (log10) as a function of the interaction distance (log10). **b)** Heatmap of the median reproducibility score (stratum adjusted correlation coefficient) across all chromosomes between replicate libraries at 50 kbp. **c)** Mean-abundance (MA) plot of the Con (Rep 1) library versus DKO (Rep 1) library before and after normalization. Plotted on the y-axis is the log-fold change of the interacting bin-pair counts between libraries and plotted on the x-axis is the log-intensity averages. **d)** Multidimensional-scaling plot (MDS) of the filtered and normalized logCPM values of bin-pairs for each library. The distance between each pair of samples was the ‘leading log fold change’, defined as the root-mean-square average of the 5000 largest log_2_ fold changes between that pair of samples.

**Supplementary Figure S3:**
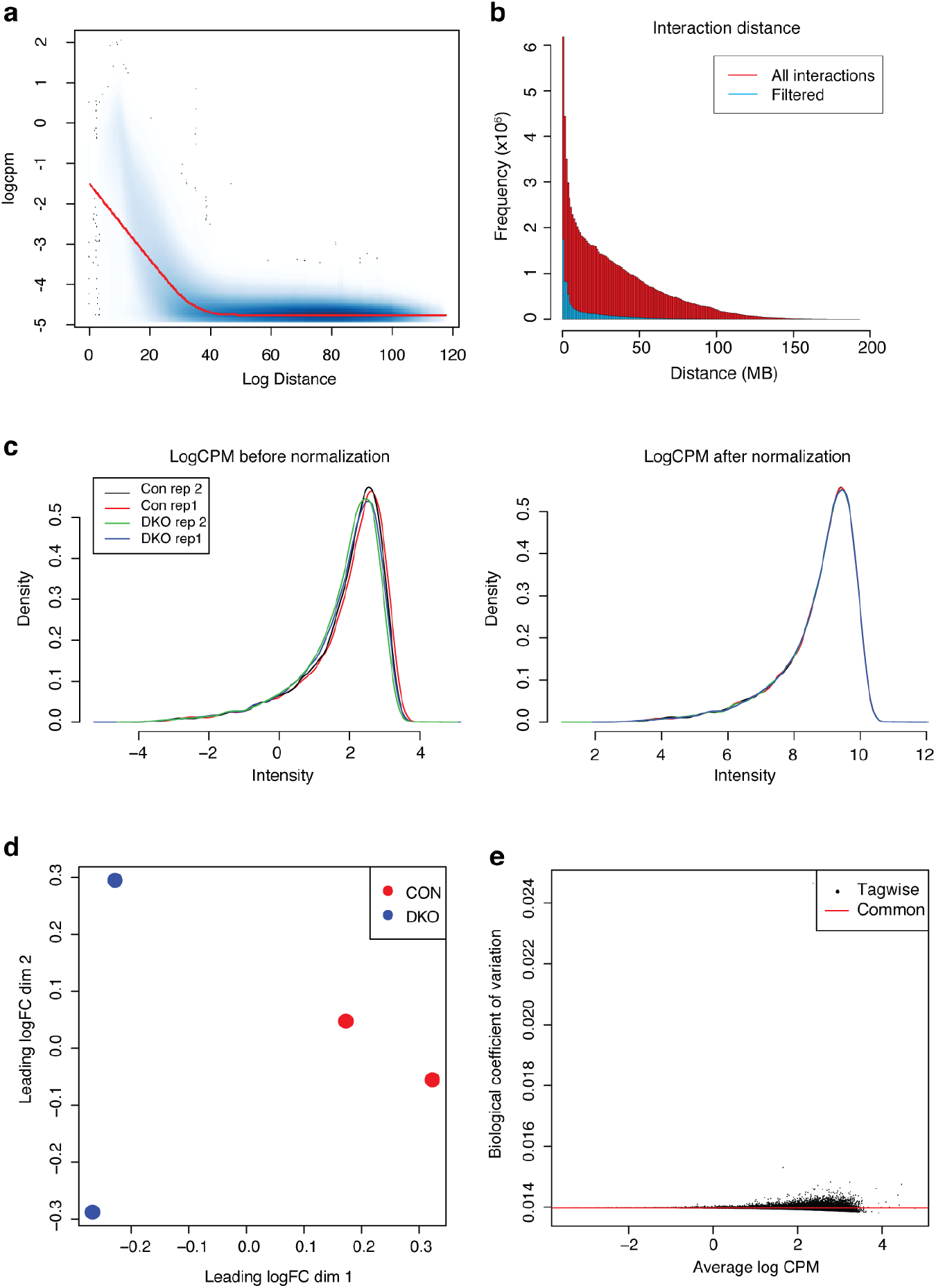
Normalization and reproducibility of gene-centric interactivity analysis (related to Figure 5). **a)** Smoothscatter plot of the average log2 counts per million (log2CPM) for all interactions as a function of distance^0.25^. A fitted loess curve in red and the minimum log2CPM value required as a function of distance in purple. **b)** Frequency of interactions at anchor distance before and after interaction filtering. **c)** LogCPM of samples before and after normalization. **d)** Multidimensional-scaling plot (MDS) of the filtered and normalized logCPM values of each promoter for each library. The distance between each pair of samples was the ‘leading log fold change’, defined as the root-mean-square average of the 500 largest log_2_ fold changes between that pair of samples. **e)** A plot of the tagwise biological coefficient of variation for each promoter versus the average log2-count per million (CPM) and common dispersion (red line). The dispersions were estimated with estimateDisp from the edgeR package with robust=TRUE and trend.method=”none”.

**Supplementary Figure S4:**
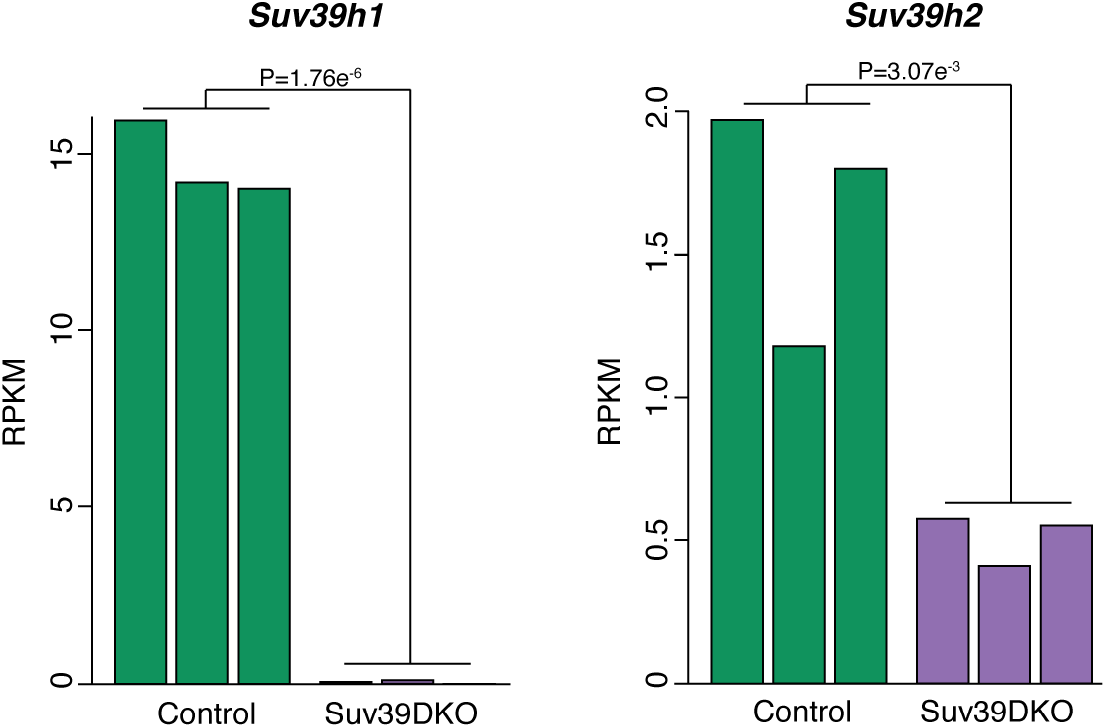
Suv39h1 and h2 expression are both significantly reduced in Suv39DKO cells (related to Figure 1). Suv39h1 and Suv39h2 mRNA expression shown as RPKM in individual RNA-seq replicates from control and Suv39DKO cells. Annotated P values shown are the adjusted p-value from *edgeR* analysis.

